# Comparative assessment of strategies to identify similar ligand-binding pockets in proteins

**DOI:** 10.1101/268565

**Authors:** Rajiv Gandhi Govindaraj, Michal Brylinski

## Abstract

**Background:** Detecting similar ligand-binding sites in globally unrelated proteins has a wide range of applications in modern drug discovery, including drug repurposing, the prediction of side effects, and drug-target interactions. Although a number of techniques to compare binding pockets have been developed, this problem still poses significant challenges.

**Results:** We evaluate the performance of three algorithms to calculate similarities between ligand-binding sites, APoc, SiteEngine, and G-LoSA. Our assessment considers not only the capabilities to identify similar pockets and to construct accurate local alignments, but also the dependence of these alignments on the sequence order. We point out certain drawbacks of previously compiled datasets, such as the inclusion of structurally similar proteins, leading to an overestimated performance. To address these issues, a rigorous procedure to prepare unbiased, high-quality benchmarking sets is proposed. Further, we conduct a comparative assessment of techniques directly aligning binding pockets to indirect strategies employing structure-based virtual screening with AutoDock Vina and rDock.

**Conclusions:** Thorough benchmarks reveal that G-LoSA offers a fairly robust overall performance, whereas the accuracy of APoc and SiteEngine is satisfactory only against easy datasets. Moreover, combining various algorithms into a meta-predictor improves the performance of existing methods to detect similar binding sites in unrelated proteins by 5-10%. All data reported in this paper are freely available at https://osf.io/6ngbs/.

## Background

Molecular functions of many proteins involve binding a variety of other molecules, including hormones, metabolites, neurotransmitters, and peptides. The analysis of ligand-protein complex structures deposited in the Protein Data Bank (PDB) [1] reveals that the majority of small organic molecules interact with specific surface regions on their macromolecular targets forming pocket-like indentations, called binding sites or binding pockets [2]. A number of computational approaches have been developed to quantify the similarity of binding sites in proteins in order to infer their molecular functions and to investigate drug-protein interactions [3]. It is now widely known that unrelated proteins may have similar binding sites with capabilities to recognize chemically similar ligands [4]. Thus, binding site matching holds a significant promise to repurpose existing drugs by facilitating the identification of novel targets. Since marketed drugs have acceptable bioavailability and safety profiles, binding site matching can efficaciously guide drug repositioning, reducing the overall costs, risks of failure, and time of drug development [5].

Accumulated evidence suggests that drugs designed for specific therapeutic targets inevitably bind to other proteins as well. Over 50% of compounds approved by the U.S. Food and Drug Administration (FDA) interact with more than five proteins often leading to unanticipated biological effects [6]. For instance, imatinib was rationally designed to treat chronic myeloid leukemia by inhibiting Bcr-Abl tyrosine-kinase [7]. Later on, imatinib was found to affect bone-resorbing osteoclasts and bone-forming osteoblasts through the inhibition of c-fms, c-kit, carbonic anhydrase II, and the platelet-derived growth factor receptor [8]; it was also shown to bind to quinone reductase 2 [9]. By employing the binding site similarity detection, off-targets can be identified at the outset of drug development in order to minimize the risk of undesired side effects.

On the other hand, the knowledge of off-binding for existing drugs opens up the possibility to find inexpensive treatments for about 7,000 rare diseases, defined as those affecting fewer than 200,000 individuals in the United States, 50,000 in Japan, and 2,000 in Australia [10]. It is estimated that only about 5% of rare diseases are of interest to the pharmaceutical industry, because developing drugs for relatively small groups of patients is considered unprofitable [11]. Compared to conventional drug discovery, much cheaper drug repositioning offers an attractive alternative to find treatments for orphan diseases [12, 13]. For instance, the rare disease late infantile neuronal ceroid lipofuscinosis (LINCL) is a neurodegenerative disorder associated with mutations in the Cln2 gene encoding tripeptidyl-peptidase I (TPP1). Because TPP1 removes tripeptides in the lysosomal compartment, its mutations lead to the accumulation of ceroid-lipofuscin causing brain cell damage. A novel use of FDA-approved lipid-lowering drugs, gemfibrozil and fenofibrate, was suggested to treat patients with LINCL by up-regulating TPP1 in brain cells [14].

Biologically meaningful similarities among proteins can be detected with sequence and structure alignment tools, such as Dynamic Programing (DP) [15], Basic Local Alignment Search Tool (BLAST) [16], TM-align [17], Combinatorial Extension (CE) [18], and Dali [19]. Although proteins with similar sequences and structures often share evolutionary ancestry and various aspects of molecular function, globally unrelated proteins may have common functional elements as well. Indeed, there are many examples of unrelated proteins binding to similar ligands and performing similar functions [4]. Despite some variations across sets of pockets interacting with the same class of small molecules [20], ligand-binding requires a certain degree of geometrical and physicochemical complementarity. Therefore, similar microenvironments generally tend to interact with similar ligands [21]. On that account, the analysis and classification of ligand-binding pockets in protein structures play an important role in drug discovery.

The last decade has witnessed a tremendous progress in the development of algorithms to measure the similarity of pockets extracted from unrelated proteins. Current methods to match binding sites can be classified into two groups, alignment-free and alignment-based techniques. Methods belonging to the former class calculate the overall similarity of binding pockets by matching various physiochemical and geometric features, and assessing the shape complementarity. For example, PocketMatch describes binding sites as lists of sorted distances encoding their shape and chemical properties, which are matched by an incremental alignment approach to compute a binding site similarity score [22]. Another algorithm, eF-seek, measures the pocket similarity according to the shapes of molecular surfaces and their electrostatic potentials [23]. Further, Patch-Surfer employs Zernike descriptors to determine the similarity of protein surfaces [24], whereas eF-site [25] and CavBase [26] capture similarities between binding sites with graph theory and clique detection algorithms.

In contrast, alignment-based methods compute local alignments of either ligand-binding residues or individual atoms in order to detect pocket similarities. Although these techniques can be computationally more expensive than alignment-free algorithms, the constructed local alignments provide valuable structural information to analyze binding modes of ligand molecules. A number of alignment-based approaches were developed to date. For instance, surface-based SiteEngine measures the pocket similarity with geometric hashing and the matching of triangles of physicochemical property centers, assuming no sequence or fold similarities [27]. SiteEngine can be applied in three modes, to scan a given functional site against a large set of complete protein structures, to compare a potential functional site with known binding sites recognizing similar features, and to search for the presence of an a priori unknown functional site in a complete protein structure. This algorithm was proposed to identify secondary binding sites of drugs that may be responsible for unwanted side effects.

The Alignment of Pockets (APoc) implements iterative dynamic programming and integer programming to calculate the optimal alignment between a pair of binding sites considering the secondary structure and fragment fitting [28]. Parameterized against millions of pocket pairs, this method can be applied not only to ligand-binding sites observed in experimental complex structures, but also to those computationally predicted by pocket-detection techniques. The Sequence Order-Independent Profile-Profile Alignment (SOIPPA) is another alignment-based approach employing a reduced representation of protein structures and sequence order-independent profile-profile alignments [29]. SOIPPA effectively detects distant evolutionary relationships despite low global sequence and structure similarities and was used to test the notion that the fold space is continuous rather than discrete. Finally, the Graph-based Local Structure Alignment (G-LoSA) has been developed to construct binding site alignments with iterative maximum clique search and fragment superimposition algorithms [30]. G-LoSA computes all possible alignments between two local structures and then the optimal solution is selected by a size-independent, chemical feature-based similarity score. Validation benchmarks demonstrated that G-LoSA efficiently identifies conserved local regions on the entire surface of a given protein. Unquestionably, these alignment-based techniques developed to predict ligand-binding sites, find template ligands, and match binding sites have a strong potential to computationally support modern drug design.

The performance of algorithms to directly compare protein binding sites heavily depends on the selection of appropriate benchmarking datasets. Several datasets have been reported to date. The Kahraman set, compiled to analyze the shapes of protein binding pockets with respect to the shapes of their ligands, contains 100 proteins binding 9 different ligands selected from different CATH homologous superfamilies [31]. Further, the Hoffman set was prepared to benchmark the performance of sup-CK, a method to quantify the similarity between binding pockets [32]. This homogeneous set contains 100 pockets extracted from non-redundant proteins binding 10 ligands of a similar size. Other datasets are composed of proteins binding a certain type of ligands. For example, the SOIPPA set comprises adenine-binding proteins as well as control proteins binding ligands that do not have the adenine moiety [29]. SOIPPA proteins represent 167 superfamilies and 146 folds according to the SCOP classification [33]. The Steroid dataset contains 8 pharmacologically relevant steroid-binding proteins complexed with 17β-estradiol, estradiol-17β-hemisuccinate, and equilenin [34]. The control subset of the Steroid dataset includes 1,854 proteins binding 334 groups of chemically diverse non-steroid molecules whose size is comparable to that of steroids. According to the SCOP classification, these target proteins represent 185 superfamilies and 150 folds.

Benchmarking datasets typically contain known binding sites extracted from the experimental structures of ligand-protein complexes, however, they may also include computationally predicted pockets. For instance, the validation of an alignment-free method to compare binding sites [35] was conducted against potential binding regions predicted with ghecom, which efficiently detects pockets on the protein surface [36]. Moreover, in addition to ligand-contacting residues detected by the Ligand-Protein Contacts (LPC) program [37], the APoc dataset contains pockets predicted with geometry-based methods LIGSITE [38] and CAVITATOR [28]. The latter was designed to be less sensitive to minor structural distortions in target structures than other techniques. Undoubtedly, employing computationally predicted pockets represents a practical approach because a number of protein structures are solved experimentally in their ligand-free conformations. On the other hand, predicted pockets with potentially incorrectly annotated binding residues certainly pose a significant challenge for binding site matching programs.

Many benchmarking sets found in the literature include structurally similar proteins as well as biologically impertinent sites that bind solvents, precipitants and additives, or are incorrectly defined based on modified amino acid residues, such as selenomethionine. These problems may cause the performance of binding site matching algorithms to be overestimated. For example, a recent paper reported that some binding sites in the APoc dataset are inadequate due to a small number of binding residues [39]. Another study revealed that although the performance of APoc against its original dataset is encouraging, it does not yield a satisfactory accuracy when applied to other datasets [30]. In contrast, G-LoSA was demonstrated to give considerably better performance than APoc in diverse benchmarks. A comprehensive review of contemporary methods to compare binding sites pointed out that these techniques generally have capabilities to predict pocket matches within diverse protein families, however, appropriate datasets to conduct an objective and unbiased assessment of their performance are lacking at present [3].

To address these issues, we carry out a thorough performance evaluation of pocket comparison algorithms. We first assess three alignment-based tools, APoc [28], SiteEngine [27] and G-LoSA [30], against an existing dataset previously compiled for that purpose. Subsequently, we construct a representative dataset comprising over one million unique pairs of drug-binding sites extracted from the PDB. The results are analyzed not only with respect to the capabilities to identify similar pockets and to construct accurate local alignments, but also taking into account the dependence of these alignments on the sequence order. Next, we propose an indirect approach to quantify the pocket similarity with structure-based virtual screening employing two popular molecular docking programs, AutoDock Vina [40] and rDock [41]. Finally, we demonstrate that combining direct and indirect approaches to compare binding sites into a metapredictor improves the accuracy of pocket matching over individual algorithms. Important aspects related to the quality of predictions made by various pocket matching tools are illustrated by representative examples. Taken together, this study sets up practical guidelines to conduct comprehensive and objective performance assessments of binding site matching programs, provides a high-quality benchmarking dataset of drug-binding pockets in globally unrelated proteins, and introduces new strategies to detect similar functional sites combining direct, alignment-based methods with indirect techniques employing structure-based virtual screening.

## Methods

### APoc dataset

The performance of binding site matching algorithms is first assessed against the APoc dataset [28], which is divided into two groups, the Subject set and the Control set. The former subset consists of non-homologous protein pairs with <30% sequence identity, binding ligands that are either identical or structurally similar at a Tanimoto coefficient [42] (TC) of ≥0.5, and sharing ≥50 atomic ligand-protein contacts of the same type. The latter subset comprises pairs of holo-proteins with <30% sequence identity, a low global structure similarity at a Template Modeling (TM)-score [43] of <0.5, and binding chemically dissimilar ligands whose TC is <0.25. TM-score ranges from 0 to 1 with values greater than 0.4 denoting statistically significant global structure similarity. Only those pockets computationally predicted by LIGSITE [38] are employed in our study. The APoc dataset comprises 34,970 Subject and 20,744 Control pairs.

### TOUGH-M1 dataset

We also compiled a new da<u>T</u>aset t<u>O</u> eval<u>U</u>ate al<u>G</u>orit<u>H</u>ms for binding site <u>M</u>atching, referred to as the TOUGH-M1 dataset, according to a procedure shown in Figure 1. First, we identified in the PDB protein chains composed of 50-999 amino acids that non-covalently bind small organic molecules (“Select ligand-bound proteins”). No constraints were imposed on the resolution to maximize the coverage and include experimentally determined structures of varied quality. Next, we retained those proteins binding a single ligand whose TC to at least one FDA-approved drug is ≥0.5 (“Select drug-like molecules”). The TC is calculated for 1024-bit molecular fingerprints with OpenBabel [44] against FDA-approved drugs in the DrugBank database [45]. Subsequently, protein sequences were clustered with CD-HIT [46] at 40% sequence similarity (“Cluster proteins”). From each homologous cluster, we selected a representative set of proteins binding chemically dissimilar ligands whose pairwise TC is <0.5 at different locations separated by at least 8 Å (“Select representative complexes”). Our intent is to evaluate the performance of binding site matching algorithms against predicted pockets. Therefore, we identified ligand-binding sites in target proteins with Fpocket 2.0, which employs Voronoi tessellation and alpha spheres to detect cavities in protein structures [47] (“Identify pockets”). We kept predicted pockets for which the Matthews correlation coefficient [48] (MCC) calculated against binding residues in the experimental complex structure reported by LPC [37] is ≥0.4. This procedure resulted in a non-redundant and representative dataset of 7,524 protein-drug complexes with computationally predicted pockets.

**Figure 1.**
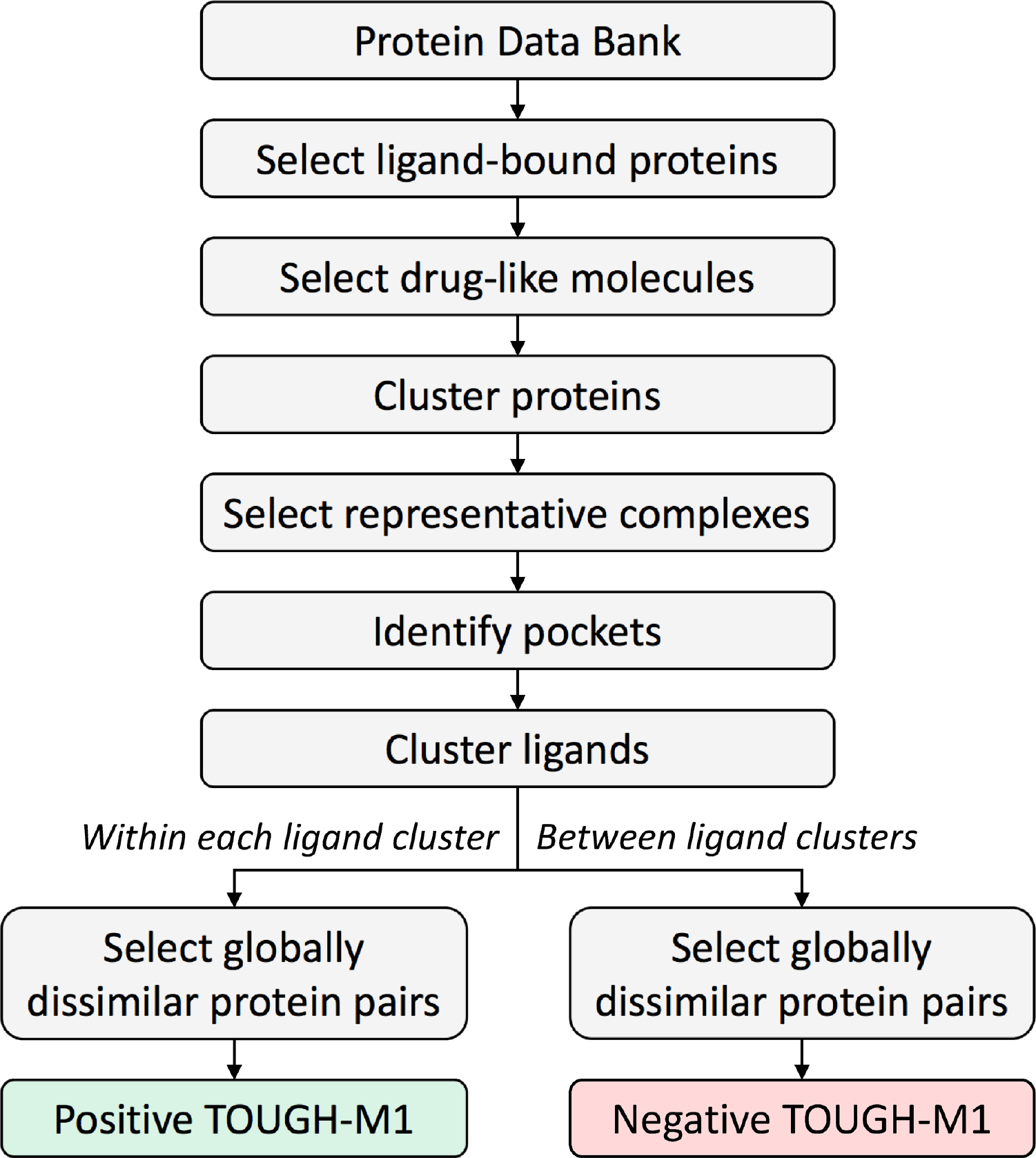
Procedure to compile the TOUGH-M1 dataset. Ligand-bound proteins selected from the Protein Data Bank are subjected to a series of filters to retain drug-like molecules and remove redundancy. Subsequently, binding pockets are computationally detected in representative complexes and the target-bound ligands are clustered to produce groups of chemically similar molecules. Finally, globally dissimilar protein pairs are identified either *within each ligand cluster* to create the Positive subset of TOUGH-M1, or *between ligand clusters* to compose the Negative subset of TOUGH-M1.

In the next step, all target-bound ligands were clustered with the SUBSET program [49] (“Cluster ligands”). Using a TC threshold of 0.7 produced 1,266 groups of chemically similar molecules. From all possible combinations of protein pairs within each cluster of similar compounds, we selected those having a TM-score of <0.4 as reported by Fr-TM-align [50] (“Select globally dissimilar protein pairs”). The Positive subset of TOUGH-M1 comprises 505,116 protein pairs having different structures, yet binding chemically similar ligands. Finally, we identified a representative structure within each group of proteins binding similar compounds, and considered all pairwise combinations of structures from different clusters that have a TM-score to one another of <0.4 (“Select globally dissimilar protein pairs”). The Negative subset of TOUGH-M1 comprises 556,810 protein pairs that have different structures and bind chemically dissimilar ligands.

### Structural comparison of binding pockets

Three algorithms to match binding sites, APoc, SiteEngine and G-LoSA, are evaluated in this study against the APoc and TOUGH-M1 datasets. APoc constructs sequence order-independent structural alignments of pockets in proteins [28]. It implements a scoring function called the Pocket Similarity (PS)-score quantifying the pocket similarity based on the backbone geometry, the orientation of side-chains, and the chemical matching of aligned pocket residues. The average PS-score for randomly selected pairs of pockets is 0.4. SiteEngine is a surface-based method developed to recognize similar functional sites in proteins having different sequences and folds [27]. The Match score is a scoring function implemented in SiteEngine to quantify the similarity of binding sites based on the number of equivalent atoms, physicochemical properties, and molecular shape complementarity. This score provides a ranking of the template sites according to the percentage of their features recognized in the target sites. Finally, we test the G-LoSA algorithm, which aligns protein binding sites in a sequence order-independent way [30]. Its scoring function, the G-LoSA Alignment (GA)-score, is calculated based on the chemical features of aligned pocket residues. The average GA-score for random pairs of local structures is 0.49. Stand-alone version of APoc v1.0b15, SiteEngine 1.0 and G-LoSA v2.1 were used in this work with default parameters for each program.

### Structure-based virtual screening

Each target binding site in the TOUGH-M1 dataset was subjected to virtual screening (VS) against a non-redundant library of 1,515 FDA-approved drugs obtained from the DrugBank database [45]. Here, the redundancy was removed with the SUBSET program [49] at a TC of 0.95. Two docking tools have been used in structure-based virtual screening, AutoDock Vina [40] and rDock [41]. Vina combines empirical and knowledge-based scoring functions with an efficient iterated local search algorithm to generate a series of docking modes ranked by the predicted binding affinity. MGL tools [51] and Open Babel [52] were used to add polar hydrogens and partial charges, as well as to convert target proteins and library compounds to the PDBQT format. For each docking ligand, the optimal search space centered on the binding site annotated with Fpocket was defined from its radius of gyration as described previously [53]. Molecular docking was carried out with AutoDock Vina 1.1.2 and the default set of parameters.

Specifically designed for high-throughput virtual screening, rDock employs a combination of stochastic and deterministic search techniques to generate low-energy ligand poses [41]. Open Babel [52] was used to convert target proteins and library compounds to the required Tripos MOL2 and SDFile formats. The docking box was defined by the rcavity program within a distance of 6 Å from the binding site center reported by Fpocket. Simulations with rDock were conducted with the default scoring function and docking parameters.

### Analysis of binding environments

The similarity of ligand-binding environments formed by two pockets is quantified with the Szymkiewicz-Simpson overlap coefficient (SSC) [54]:

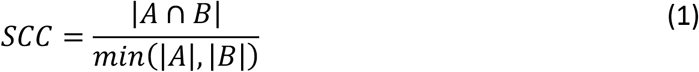

where *A* and *B* are the lists of protein-ligand contacts within the two pockets according to the LPC program [37]. In order to calculate the intersection, we employ a similar procedure to that used to compile the APoc dataset [28]. Specifically, two contacts in different structures are of the same type if the ligand atoms are equivalent according to the chemical alignment by the kcombu program [55] and the protein residues belong to the same group (I-VIII) defined as: I (LVIMC), II (AG), III (ST), IV (P), V (FYW), VI (EDNQ), VII (KR), VIII (H) [56].

### Evaluation metrics

Recognizing those pockets binding similar ligands in different proteins is essentially a binary classification problem, *viz.* pairs of pockets are classified as either similar or dissimilar. Therefore, the performance of pocket matching algorithms can be evaluated with the Receiver Operating Characteristic (ROC) analysis and the corresponding area under the ROC curve (AUC). Pairs of pockets binding either the same or chemically similar ligands in the APoc dataset (Subject) and the TOUGH-M1 dataset (Positive) are positives, *P*, whereas those pockets binding dissimilar ligands, the Control subset of the APoc dataset and Negative subset of TOUGH-M1 are negatives, *N*. ROC analysis is based on a true positive rate (*TPR*) also known as the sensitivity, and a false positive rate (*FPR*) also known as the fall-out, defined as:

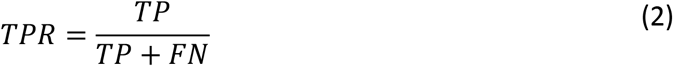

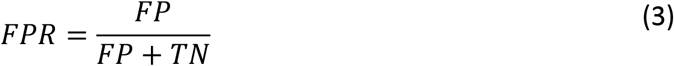

where *TP* is the number of true positives, i.e. Subject (APoc) and Positive (TOUGH-M1) pairs classified as similar pockets, and *TN* is the number of true negatives, i.e. Control (APoc) and Negative (TOUGH-M1) pairs classified as dissimilar pockets. *FP* is the number of false positives or over-predictions, i.e. Control (APoc) and Negative (TOUGH-M1) pairs classified as similar pockets, and *FN* is the number of false negatives or under-predictions, i.e. Subject (APoc) and Positive (TOUGH-M1) pairs classified as dissimilar pockets.

The sequence order of alignments constructed by individual binding site matching algorithms is quantified by the Kendall *τ* rank correlation coefficient [57]. The Kendall *τ* measuring the degree of the ordinal association of binding site residues is given by:

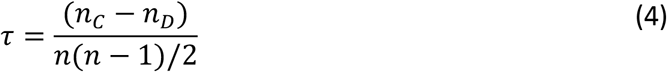

where *n*_*c*_ and *n*_*D*_ are the numbers of concordant and discordant pairs, respectively. A pair of pocket residues is concordant if their order in the protein sequence and the local alignment is the same, otherwise the residue pair is discordant. Sequence order-dependent alignments tend to have high Kendall *τ* values, with *τ* = 1 for completely sequential alignments calculated by e.g. DP [15], BLAST [16], TM-align [17], CE [18], and DALI [19]. A Kendall *τ* value of -1 corresponds to an alignment in which the order of one binding site is reversed. Fully sequence order-independent alignments theoretically have the Kendall *τ* of around 0.

In addition to direct pocket matching, the pocket similarity across the TOUGH-M1 dataset is also indirectly measured with structure-based virtual screening. Here, the statistical dependence between the ranking of library compounds against a pair of target pockets is evaluated by non-parametric Spearman’s ρ rank correlation coefficient:

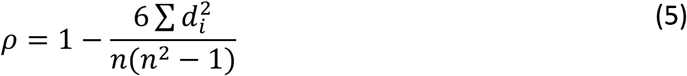

where *d*_i_ is a difference between the ranks of a compound docked against two target pockets and *n* is the total number of screening compounds, in our case *n* = 1,515 (the number of FDA-approved drugs). Spearman’s ρ ranges from -1 to 1 with negative and positive values corresponding to an indirect and direct correlation, respectively. Therefore, a high and positive Spearman’s ρ indicates that two target binding sites are chemically similar, i.e. tend to bind similar compounds.

Finally, the quality of local alignments is assessed by a ligand root-mean-square deviation (RMSD) calculated upon the superposition of binding pockets residues. Here, the assumption is that the correct alignment of binding residues causes bound ligands to adopt a similar orientation. On that account, two proteins are first superposed using Cα atoms of equivalent binding residues according to a given local alignment and then the RMSD is calculated over ligand heavy atoms. In addition, the accuracy of pocket alignments is evaluated with the MCC against reference alignments obtained by the superposition of bound ligands [34].

## Results and Discussion

### Performance of pocket matching algorithms on the APoc dataset

We begin by evaluating the performance of APoc, SiteEngine, and G-LoSA on the APoc dataset [28] with pockets predicted by LIGSITE [38]. The solid gray line in Figure 2 shows the accuracy of predicted ligand-binding pockets assessed by the MCC against experimental binding residues reported by LPC [37]. The majority of pockets are accurately predicted with the MCC of ≥0.6 (≥0.4) in 44% (81.7%) of the cases. Further, as many as 88.3% of the best pockets are top-ranked, corresponding to the largest cavity detected in a given target structure (inset in Figure 2, gray bars).

**Figure 2.**
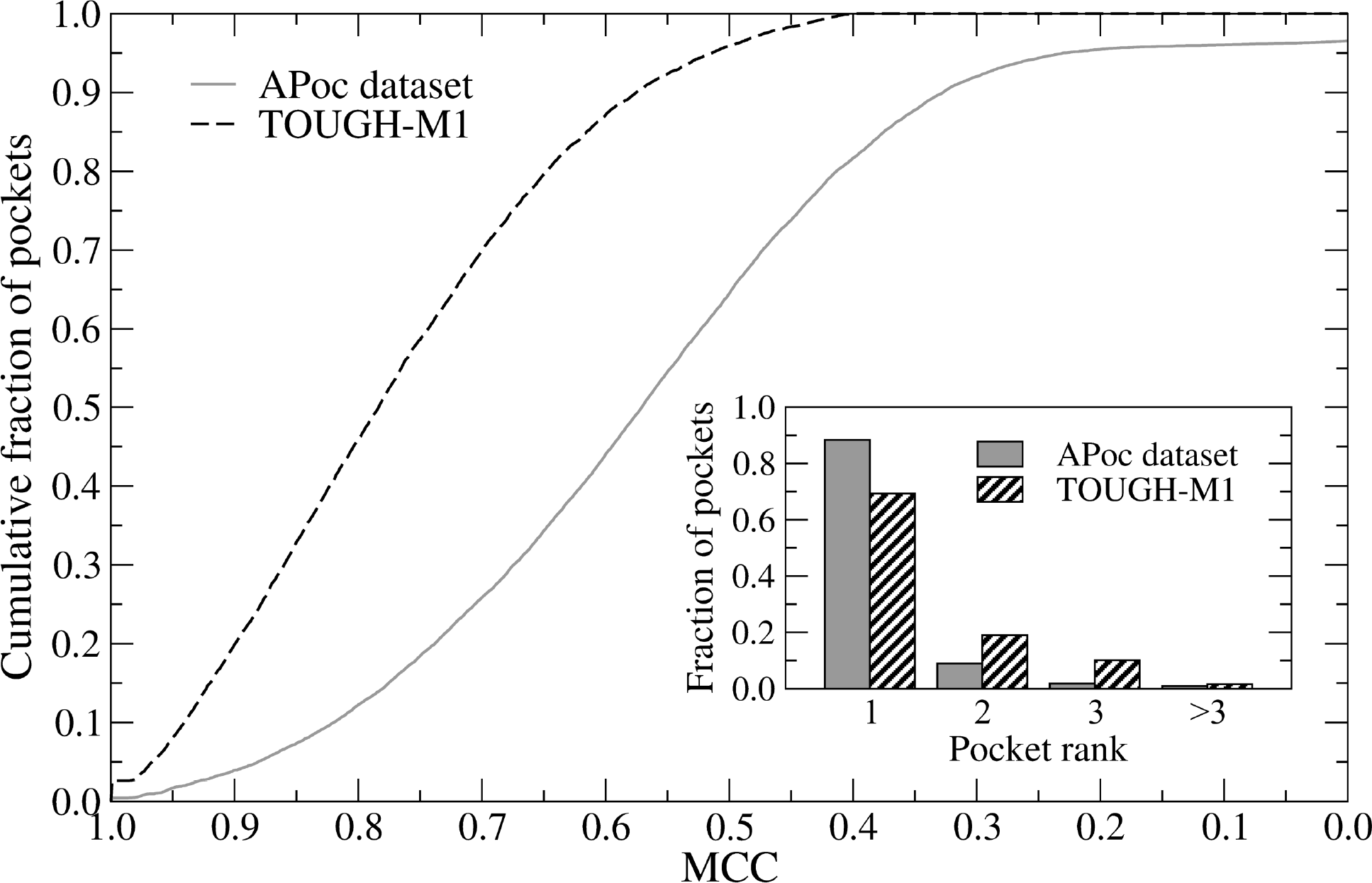
Ligand-binding pockets predicted across APoc and TOUGH-M1 datasets. The accuracy of pocket identification is evaluated by the Matthews correlation coefficient (MCC) calculated over binding residues against experimental complex structures. Inset: Pocket ranking assessed by the fraction of targets, for which the best pocket is found at a particular rank shown on the *x*-axis.

A ROC analysis is conducted to assess the performance of APoc, SiteEngine, and G-LoSA detecting similar binding pockets for the APoc dataset (Figure 3 and Table 1). Here, APoc and G-LoSA are the best performing algorithms with AUC values of 0.82 and 0.77, respectively, whereas the AUC for SiteEngine is somewhat lower (0.60). We also compare the performance of pocket matching tools to that obtained using the global sequence identity computed with DP [15] as well as the global structure similarity calculated with Fr-TM-align [50]. Distinguishing between proteins binding similar and dissimilar compounds on the basis of just the sequence identity yields an AUC of 0.56, which is fairly close to the performance of a random classifier. This is expected because a sequence similarity threshold of 30% between the two associated proteins was used to compile both the Subject (pockets binding similar ligands) and the Control (pockets binding dissimilar ligands) sets [28]. Nonetheless, classifying pockets based on the global structure similarity of the associated proteins measured by the TM-score [43] yields a much higher performance on this dataset with an AUC of 0.75. This can be expected as well because a global TM-score threshold of 0.5 was imposed only on target pairs within the Control set and not on those within the Subject set [28].

**Figure 3.**
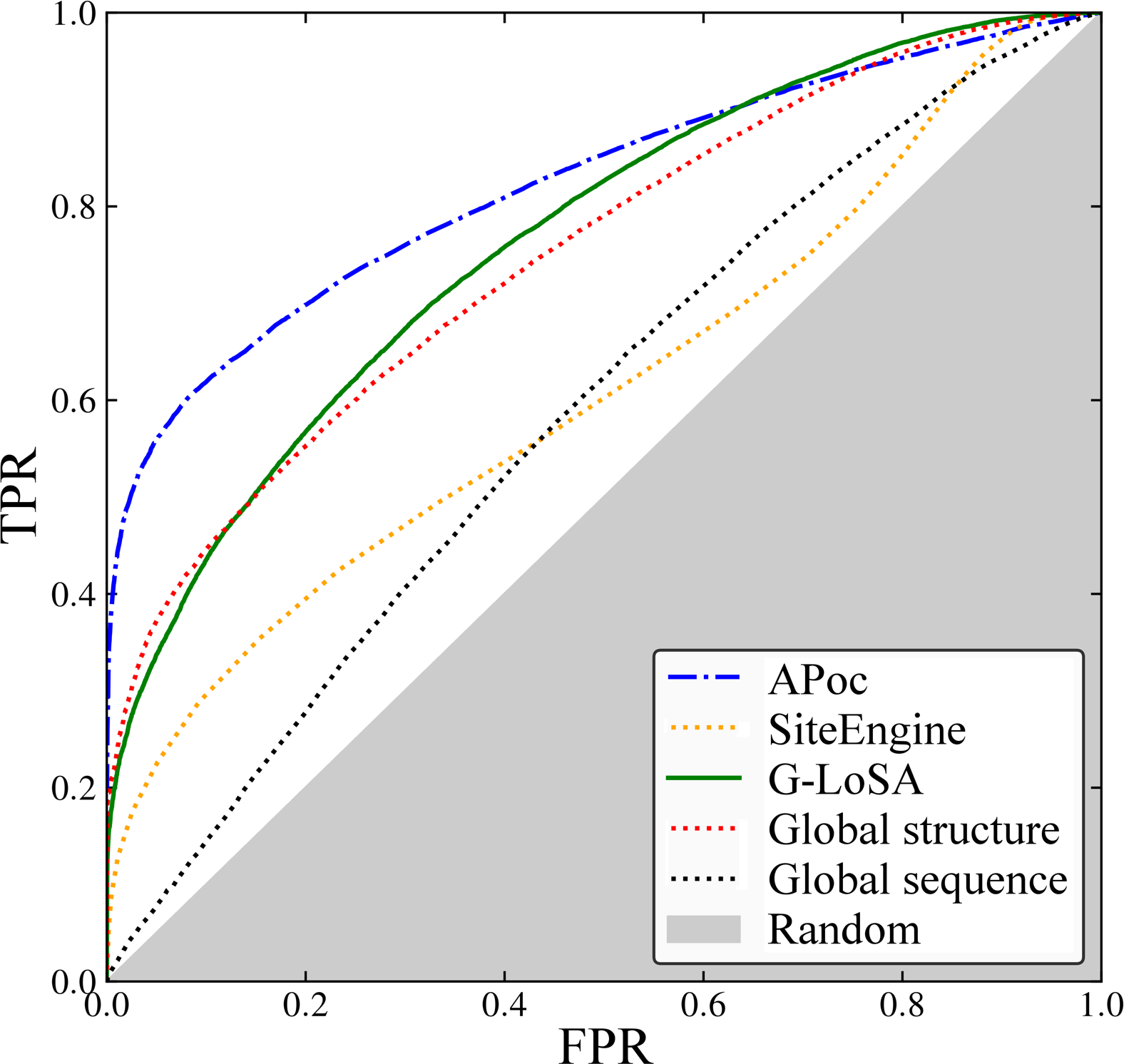
ROC plot assessing the performance of APoc on the APoc dataset. The dashed-dotted blue line shows the performance against the entire dataset as reported in the original publication of APoc. The dotted red line evaluates to the performance for globally similar Subject pairs, whereas the solid green line corresponds to the performance of APoc when globally similar Subject pairs are excluded from benchmarks. The *x*-axis shows the false positive rate (FPR) and the *y*-axis shows the true positive rate (TPR). The gray area represents a random prediction.

**Table 1.**
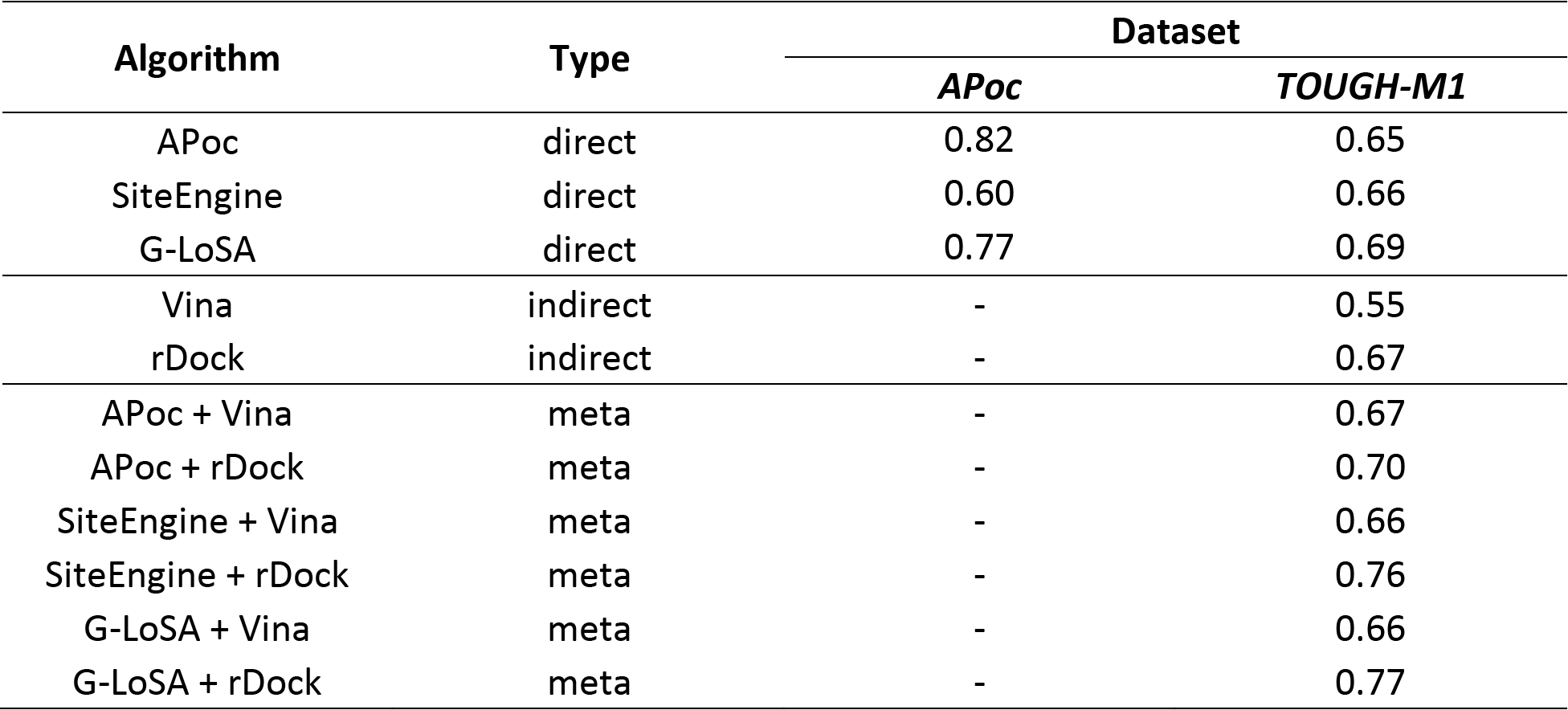
Performance of various strategies to match ligand-binding pockets assessed with the area under the ROC curve (AUC) Three types of algorithms are evaluated, direct methods based on the local alignment between a pair of pockets, indirect techniques employing structure-based virtual screening to detect chemically similar binding sites, and meta-predictors combining a direct and an indirect algorithm. Direct approaches are benchmarked on APoc and TOUGH-M1 datasets, whereas indirect methods and meta-predictors are assessed against the TOUGH-M1 dataset only.

To further assess how the performance of pocket matching algorithms is affected by the global structure similarity, we identified globally structurally similar protein pairs included in the APoc dataset. Only 4.7% Control pairs have a TM-score of ≥0.4, whereas as many as 36.5% Subject pairs are structurally similar at a TM-score of ≥0.4. On that account, we divide the Subject set into two subsets, one comprising 12,773 pairs having globally similar structures and the other consisting of 22,197 pairs with different global structures. Figure 4 shows a ROC plot assessing the performance of APoc, set as an example, against the entire dataset (the dashed-dotted blue line), as well as globally similar Subject pairs (the dotted red line) and dissimilar Subject pairs (the solid green line). Note that we employ the entire Control set in this analysis because it contains only a negligible fraction of structurally similar proteins. On the entire dataset, APoc achieves the sensitivity values of 43.6% and 55.9% at an FPR of 1% and 5%, respectively, just as reported in the original publication [28]. However, the sensitivity values are as high as 78.2% at 1% FPR and 87.4% at 5% FPR when only globally similar Subject pairs are included in the ROC analysis. Furthermore, excluding any global structure similarity from the benchmarking dataset dramatically decreases the performance of APoc to the sensitivity values of only 23.8% and 37.9% at an FPR of 1% and 5%, respectively. In the following sections, we look into the potential causes of this high discrepancy in the performance of APoc and the correspondingly uneven results.

**Figure 4.**
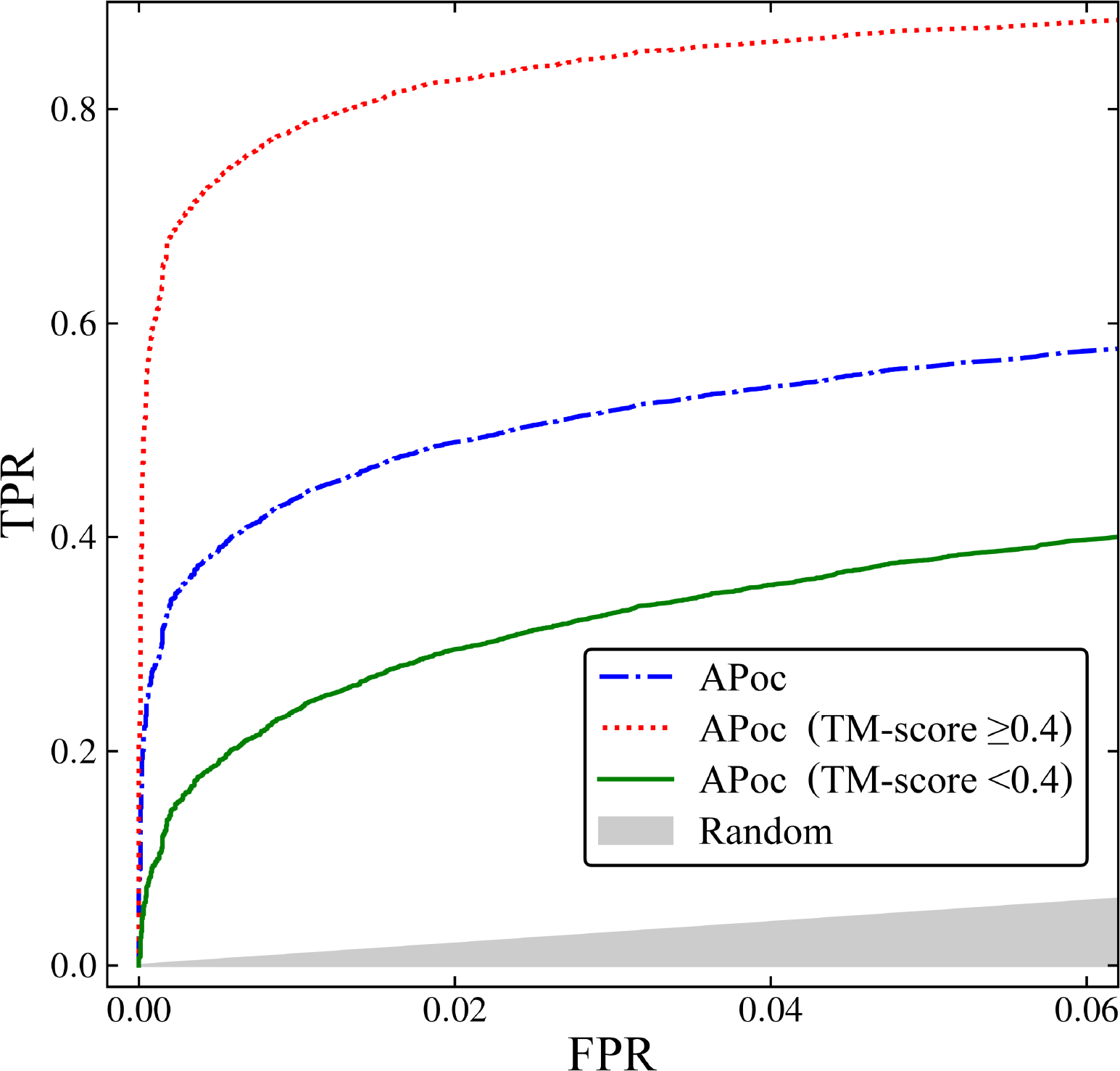
ROC plot assessing the performance of APoc on the APoc dataset. The dashed-dotted blue line shows the performance against the entire dataset as reported in the original publication of APoc. The dotted red line evaluates to the performance for globally similar Subject pairs, whereas the solid green line corresponds to the performance of APoc when globally similar Subject pairs are excluded from benchmarks. The *x*-axis shows the false positive rate (FPR) and the *y*-axis shows the true positive rate (TPR). The gray area represents a random prediction.

### Characteristics of alignments constructed for the APoc dataset

We found that local alignment scores reported by pocket matching algorithms, APoc in particular, are correlated with the global structure similarity. Figure 5 shows that the Pearson correlation coefficient (PCC) between the TM-score and the PS-score from APoc (Figure 5A), the Match score from SiteEngine (Figure 5B), and the GA-score from G-LoSA (Figure 5C) calculated across the Subject set are 0.72, 0.53, and 0.52, respectively. Because significantly fewer structurally similar proteins are included in the Control set, there is no correlation between the global and local structure similarity for SiteEngine (PCC = 0.10) and G-LoSA (PCC = -0.02), however, PS-score values computed by APoc still correlate with the TM-score (PCC = 0.48).

**Figure 5.**
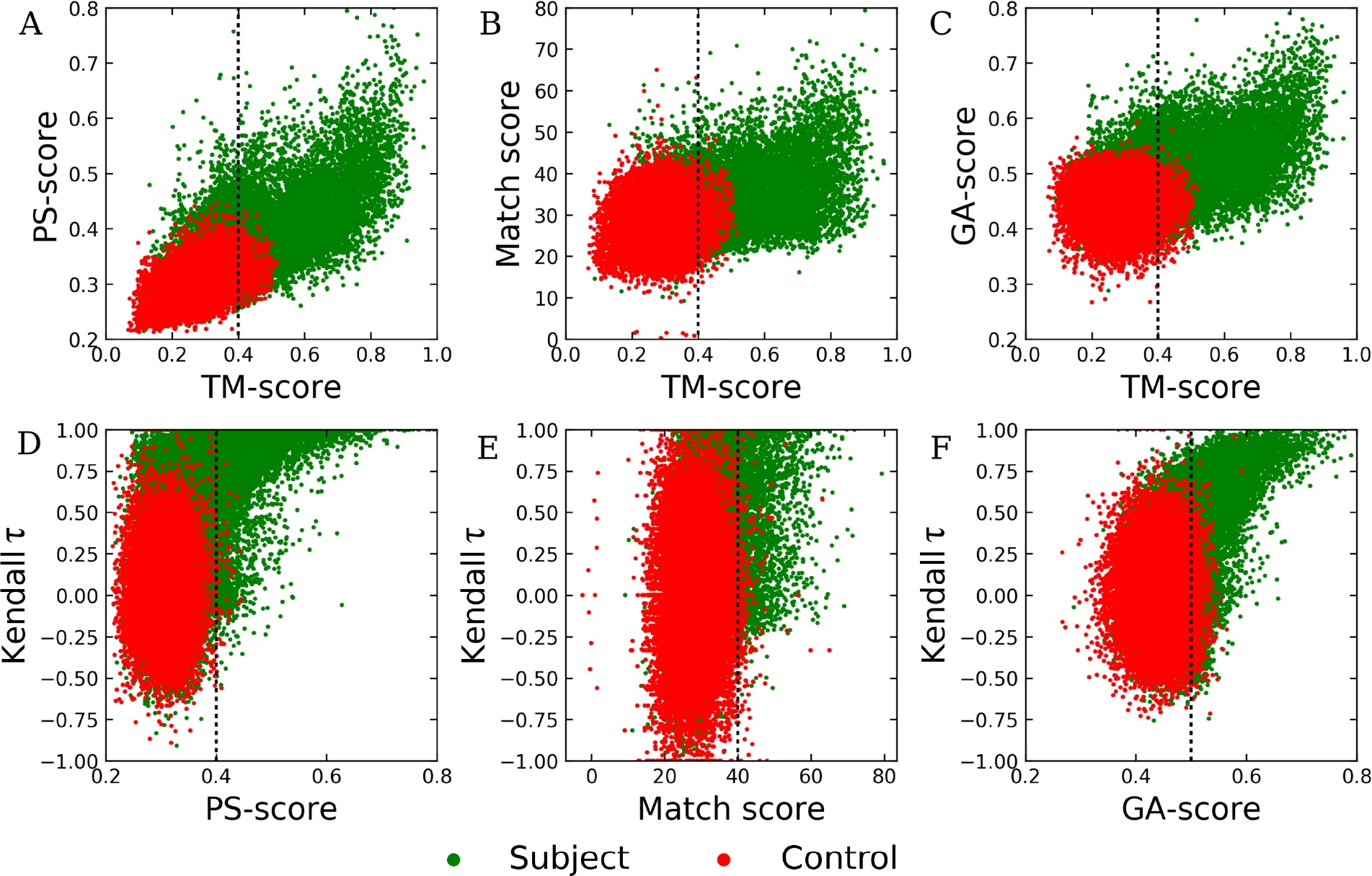
Characteristics of local alignments constructed by pocket matching algorithms for the APoc dataset. Top panel shows the correlation between the global structure similarity score, TM-score, and the local alignment score, (**A**) PS-score by APoc, (**B**) Match score by SiteEngine, and (**C**) GA-score by G-LoSA. Bottom panel shows the correlation between local alignment scores reported by individual pocket matching algorithms, (**D**) APoc, (**E**) SiteEngine, and (**F**) G-LoSA, and Kendall *τ* measuring the degree of the ordinal association of binding site alignments. Dotted lines mark thresholds for statistically significant alignments.

Another important issue that needs to be addressed is whether binding site similarity scores reported by pocket matching algorithms depend on the sequence order of the constructed alignments. Kendall *τ* values measuring the ordinal association of binding residues in local alignments are broadly distributed across the Control set with a central tendency around 0 (red dots in Figures 5D and 5E). Clearly, pocket matching algorithms tested in this study construct fully sequence order-independent alignments for dissimilar pockets extracted from unrelated protein structures, which are also assigned low similarity scores. Nevertheless, the distribution of green dots in Figure 5D shows that the vast majority of highly significant PS-score values computed by APoc for the Subject set actually correspond to mainly sequential alignments as indicated by high Kendall *τ* values. This is also the case for G-LoSA (Figure 5F) and, to a lesser extent, for SiteEngine (Figure 5E). The majority of similar pockets extracted from unrelated structures in the Subject set are assigned low similarity scores that are no different from the Control set. It appears that high pocket similarity scores are computed for mainly sequential local alignments constructed for those target pairs in the Subject set having globally similar structures. Considering that 36.5% Subject pairs are structurally similar, these results corroborate ROC plots presented in Figure 4.

### Representative examples from the APoc dataset

We selected a couple of representative examples to illustrate difficulties in conducting an objective assessment of the performance of pocket matching algorithms on the APoc dataset. The first case is a pair of protein kinases, inositol 1,4,5-trisphosphate 3-kinase B from mouse bound to ATP [IP(3)-3KB, PDB ID: 2aqx, chain A, 287 aa) [58] and inositol polyphosphate multikinase 2 from yeast bound to ADP (Ipk2, PDB ID: 2if8, chain A, 255 aa) [59]. Binding sites in both targets are highly similar with a SSC of 0.67. Figure 6 shows local alignments constructed for IP(3)-3KB (violet) and Ipk2 (blue). The reference alignment presented in Figure 6A was obtained by superposing proteins using the coordinates of bound ligands. Because adenine nucleotides bound to these targets have slightly different internal conformations, the ligand RMSD in the reference alignment is 0.81 Å. The alignment by G-LoSA (Figure 6B) has a GA-score of 0.59, revealing a significant similarity of both binding sites. Likewise, the PS-score reported by APoc for these target pockets is 0.57 with the corresponding *p*-value of 1.08 ×10^−6^ (Figure 6C), also indicating a significant pocket similarity.

**Figure 6.**
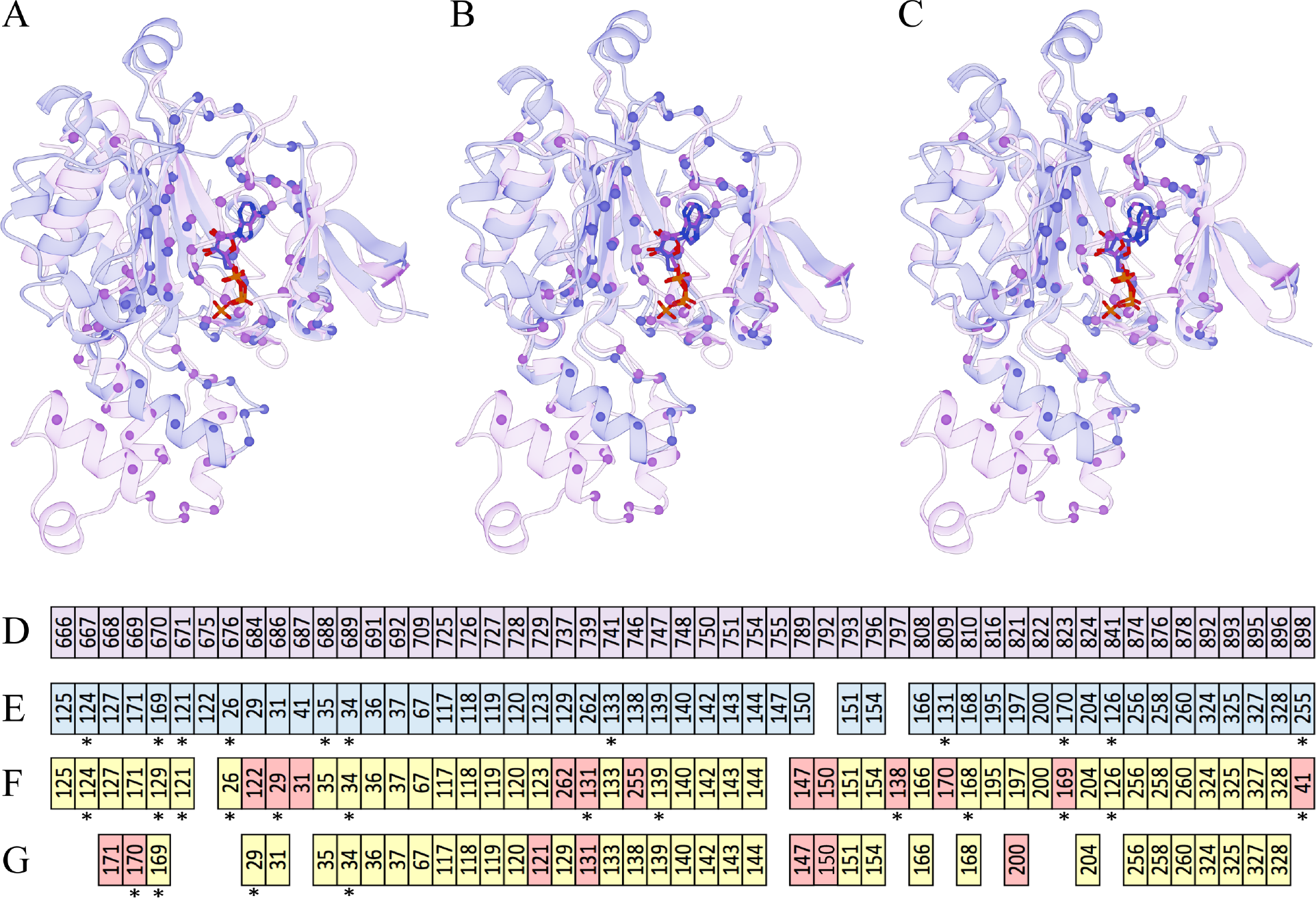
Examples of local alignments constructed by pocket matching algorithms for a pair of structurally similar ATP/ADP-binding proteins in the APoc dataset. Inositol 1,4,5-trisphosphate 3-kinase B [IP(3)-3KB, violet] is aligned to inositol polyphosphate multikinase 2 (Ipk2, blue). (A) The reference alignment obtained by the superposition of bound ligands is compared to pocket alignments by (**B**) G-LoSA and (**C**) APoc. Cα atoms of binding residues are represented by solid spheres, whereas adenine nucleotides are shown as solid sticks. (**D-G**) Textual pocket alignments between IP(3)-3KB and Ipk2, (**D↔E**) the reference alignment and those constructed by (**D↔F**) G-LoSA and (**D↔G**) APoc. Each box represents a binding residue. Alignments are sorted by the sequence order of IP(3)-3KB in **D** (violet). Equivalent residues in Ipk2 are colored in blue in E, whereas in **F** and **G**, correctly aligned residues with respect to the reference alignment are colored in yellow and misaligned residues are colored in red. Alignment positions reversing the sequence order are marked by asterisks.

The alignment accuracy can be assessed by an RMSD computed over ligand heavy atoms upon the superposition of aligned binding residues. Both programs constructed a correct alignment with a ligand RMSD of 1.00 Å (G-LoSA) and 0.95 Å (APoc). In addition, textual alignments between IP(3)-3KB and Ipk2 are shown in Figures 6D-G. The reference alignment (Figure 6D↔E) is mostly sequential with the Kendall *τ* of 0.67. MCC values calculated for alignments generated by G-LoSA (Figure 6D↔F) and APoc (Figure 6D↔G) are as high as 0.74 and 0.71, respectively. Despite a low sequence identity of 23.1%, both kinases are structurally globally similar with a TM-score of 0.74 and a Cα-RMSD of 1.93 Å over 204 aligned residues. Consequently, pocket alignments are mainly sequential with the Kendall *τ* of 0.62 for G-LoSA and 0.77 for APoc. These high ordinal association values are likely the reason for significant pocket similarity scores because these quantities depend on each other as shown in Figures 5D and 5F.

To further look into this issue, we consider another ADP-bound protein, glutathione synthetase from *Escherichia coli* (GSHase, PDB ID: 1gsa, chain A, 314 aa) [60], whose sequence identity to IP(3)-3KB is 19.9%. Although GSHase and IP(3)-3KB are structurally unrelated with a TM-score of 0.28 and a Cα-RMSD of 5.56 Å over 130 aligned residues, their binding sites are similar with a SSC of 0.47. Figure 7 shows local alignments between IP(3)-3KB (violet) and GSHase (green). The ligand RMSD in the reference alignment presented in Figure 7A is 1.10 Å. Adenine nucleotides bound to IP(3)-3KB and GSHase adopt a similar orientation with an RMSD of 1.28 Å when target proteins are superposed according to the alignment by G-LoSA (Figure 7B). Equally important, the pocket alignment by G-LoSA is assigned a significant similarity score of 0.51. In contrast, the similarity of ATP/ADP-binding sites in IP(3)-3KB and GSHase was not recognized by APoc, which assigned a low PS-score of 0.32 and an insignificant *p*-value of 0.33 to this pair of target pockets. Further, the ligand RMSD calculated for the alignment reported by APoc is as high as 14.21 Å indicating that equivalent binding residues have not been correctly identified (Figure 7C).

**Figure 7.**
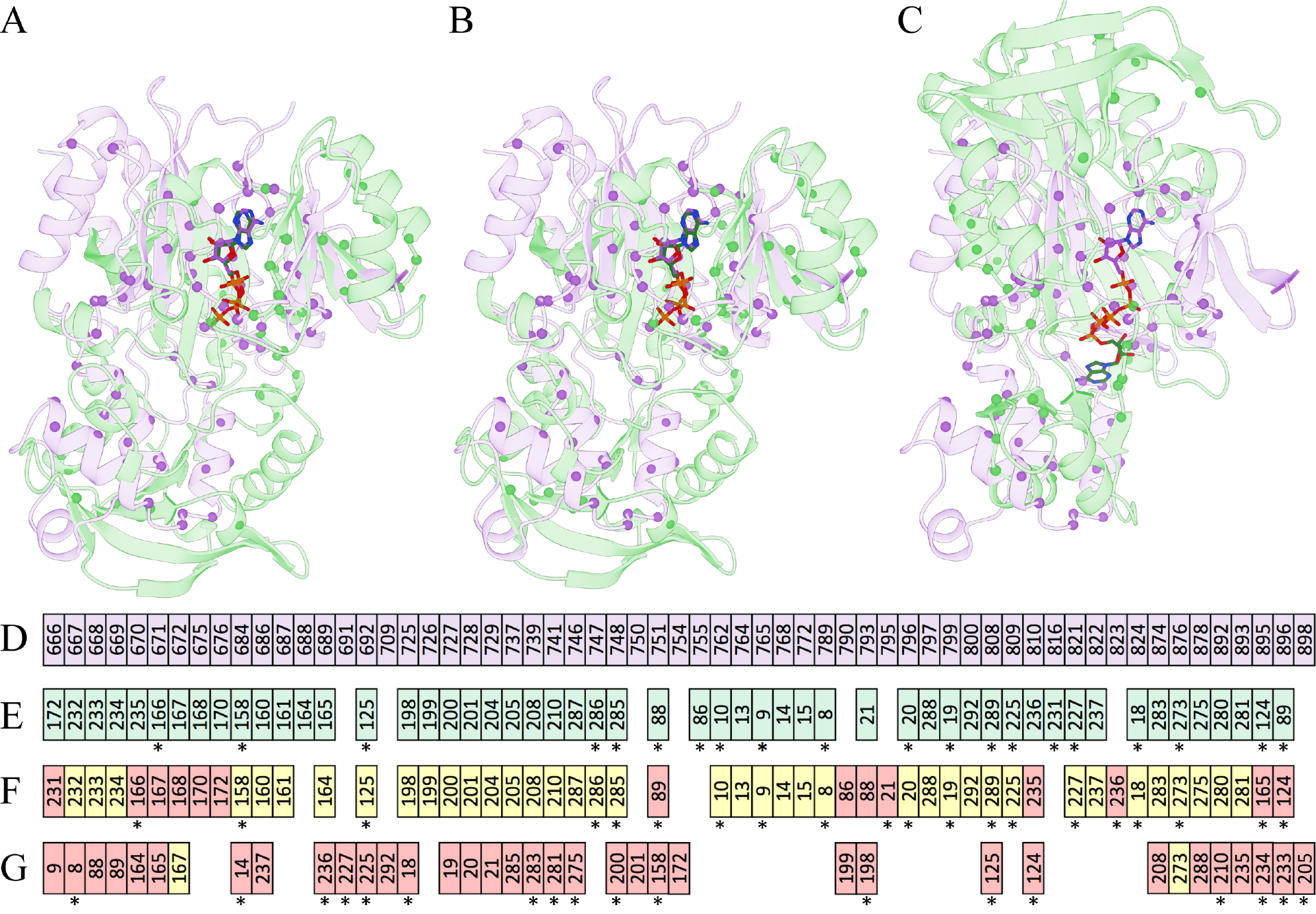
Examples of local alignments constructed by pocket matching algorithms for a pair of structurally dissimilar ATP/ADP-binding proteins in the APoc dataset. Inositol 1,4,5-trisphosphate 3-kinase B [IP(3)-3KB, violet] is aligned to glutathione synthetase (GSHase, green). (**A**) The reference alignment obtained by the superposition of bound ligands is compared to pocket alignments by (**B**) G-LoSA and (**C**) APoc. Cα atoms of binding residues are represented by solid spheres, whereas adenine nucleotides are shown as solid sticks. (**D-G**) Textual pocket alignments between IP(3)-3KB and GSHase, (**D↔E**) the reference alignment and those constructed by (**D↔F**) G-LoSA and (**D↔G**) APoc. Each box represents a binding residue. Alignments are sorted by the sequence order of IP(3)-3KB in *D* (violet). Equivalent residues in Ipk2 are colored in green in **E**, whereas in **F** and **G**, correctly aligned residues with respect to the reference alignment are colored in yellow and misaligned residues are colored in red. Alignment positions reversing the sequence order are marked by asterisks.

These results are further corroborated by textual pocket alignments between IP(3)-3KB and GSHase shown in Figures 7D↔E (the reference alignment), 7D↔F (G-LoSA), and 7D↔G (APoc). Owing to the fact that target structures are globally unrelated, the reference alignment is fully sequence-order-independent with the Kendall *τ* of 0.09. Encouragingly, the alignment by G-LoSA is not only non-sequential with the Kendall *τ* of 0.10, but it is also highly accurate as assessed by an MCC of 0.71 against the reference alignment. On the other hand, APoc constructed an inaccurate (an MCC of 0.03) and partially sequential (the Kendall *τ* of 0.29) local alignment between IP(3)-3KB and GSHase. These case studies illustrate difficulties in conducting an objective evaluation of pocket matching algorithms on the APoc dataset containing a significant number of structurally similar Subject pairs. Specifically, high similarity scores are typically computed by APoc from sequential alignments constructed for those targets having globally similar structures, whereas its performance against structurally unrelated target pairs is notably lower.

### TOUGH-M1 dataset

In order to factor out the correlation between the local and global structure similarity as well as the dependence of the similarity score on the sequence order of target proteins, we compiled a new set of over 1 million pocket pairs, the TOUGH-M1 dataset. TOUGH-M1 is not only non-redundant and representative, but it comprises pairs of pockets extracted from proteins with globally unrelated sequences and structures. It is further divided into two subsets, pairs of pockets binding chemically similar molecules (Positive) and pairs of pockets binding different ligands (Negative). Because we consider pockets predicted by a geometrical approach, benchmarking results against the TOUGH-M1 dataset are relevant for the subsequent large-scale applications utilizing computationally predicted binding sites. The dashed black line in Figure 2 shows that the MCC calculated against experimental binding residues is ≥0.4 for all pockets, which was one of the criteria to construct this dataset, and it is ≥0.6 for as many as 87.1% predicted pockets. Therefore, pockets in the TOUGH-M1 dataset are generally more accurate than those predicted by LIGSITE in the APoc dataset [28]. In terms of the pocket ranking, 69.3% pockets correspond to the largest, top-ranked cavity identified by Fpocket (inset in Figure 2, striped bars).

We also analyze the similarity of binding environments for pairs of pockets in the TOUGH-M1 dataset with the contact-based overlap coefficient, SCC. The distribution of SCC values across the Positive and Negative subsets of TOUGH-M1 are shown in Figure 8. Because pairs of pockets in the Positive subset bind either exactly the same compound or at least chemically very similar molecules, a significant overlap between their binding environments can be observed with a median SCC of 0.30. This result is in line with other studies demonstrating that although the complete functional sites binding similar ligands may be somewhat different as a result of binding to compounds with different compositions or conformations, they share similar subsites interacting with similar ligand fragments [29]. Further, the high similarity of binding environments across the Positive pairs likely arises from the presence of strongly conserved anchor functional groups in ligands binding to common binding sites within evolutionarily related, but distant protein families [61]. In contrast, binding environments formed by pockets in the Negative set are very different from one another with a median SCC of only 0.05. On that account, algorithms designed to recognize similar pockets should be able to effectively distinguish between Positive and Negative pairs even in the absence of any global structure similarity.

**Figure 8.**
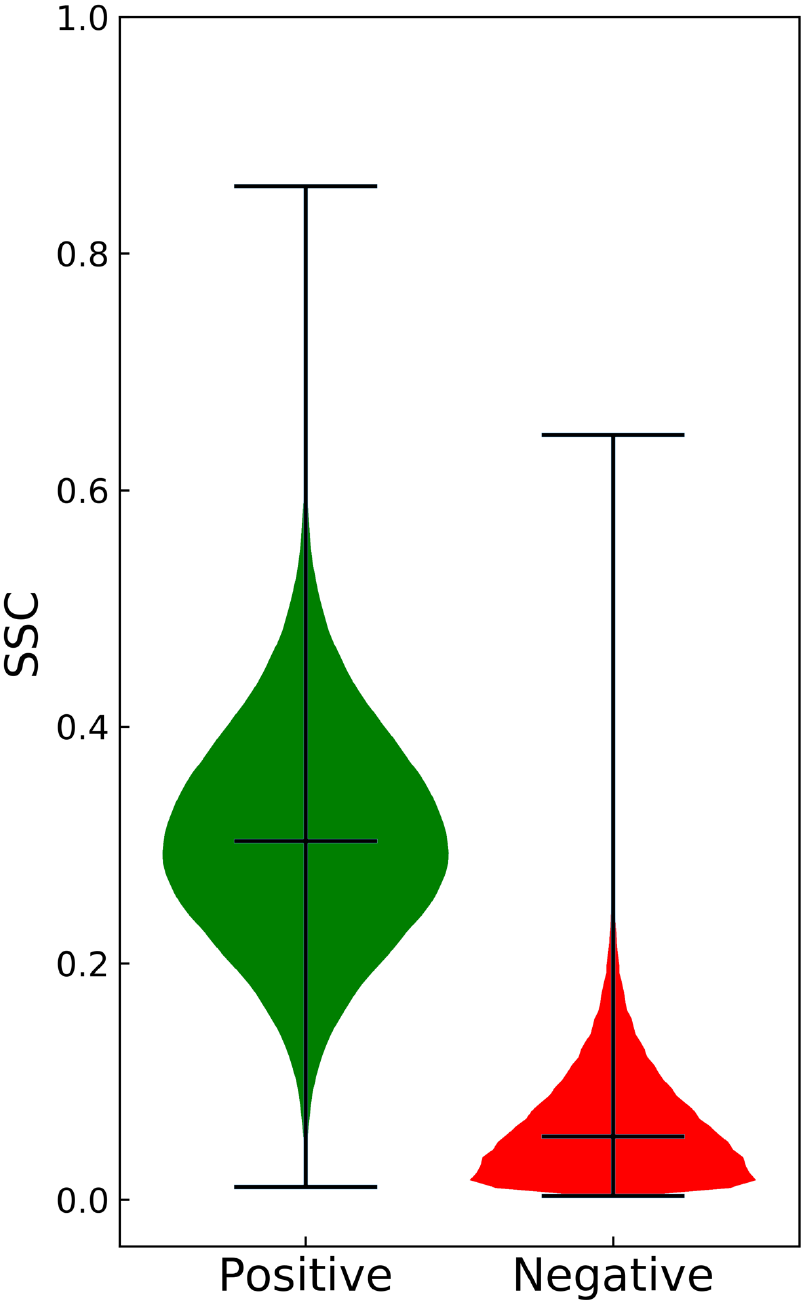
Similarity of binding environments across the Positive and Negative subsets of the TOUGH-M1 dataset. The overlap of atomic ligand-protein contacts of a similar chemical type is measured with the Szymkiewicz-Simpson coefficient (SCC). Three black horizontal lines in each group represent (from the top) the maximum value, the median, and the minimum value.

### Performance of pocket matching algorithms on the TOUGH-M1 dataset

The performance of binding site alignment algorithms in identifying similar binding sites across the TOUGH-M1 dataset is assessed with the ROC analysis. Results for APoc, SiteEngine, and G-LoSA are shown in Figure 9. Notably, the performance of APoc with an AUC of 0.65 is lower than SiteEngine and G-LoSA, which yield AUC values of 0.66 and 0.69, respectively. Further, the ROC analysis includes the pairwise global sequence identity computed with DP [15] and global structure similarity calculated with Fr-TM-align [50]. As expected, the classification of pockets based on sequence identity gives an AUC of 0.55, which is close to that of a random classifier. In contrast to the APoc dataset, the TM-score [43] yields a very low performance with an AUC of 0.51, therefore, Positive and Negative protein pairs in the TOUGH-M1 dataset cannot be separated simply based on the similarity of their global structures. This result is not surprising because TOUGH-M1 was compiled at a global sequence similarity threshold of 40% and a TM-score threshold of <0.4. Our benchmarking calculations reveal that G-LoSA is the only pocket matching algorithm offering a fairly robust performance on both datasets with AUC values of 0.77 (APoc) and 0.69 (TOUGH-M1). This robustness likely develops from an efficient search algorithm, which generates sequence order-independent alignments by solving the assignment problem with a combination of an iterative maximum clique search and a fragment superimposition [30].

**Figure 9.**
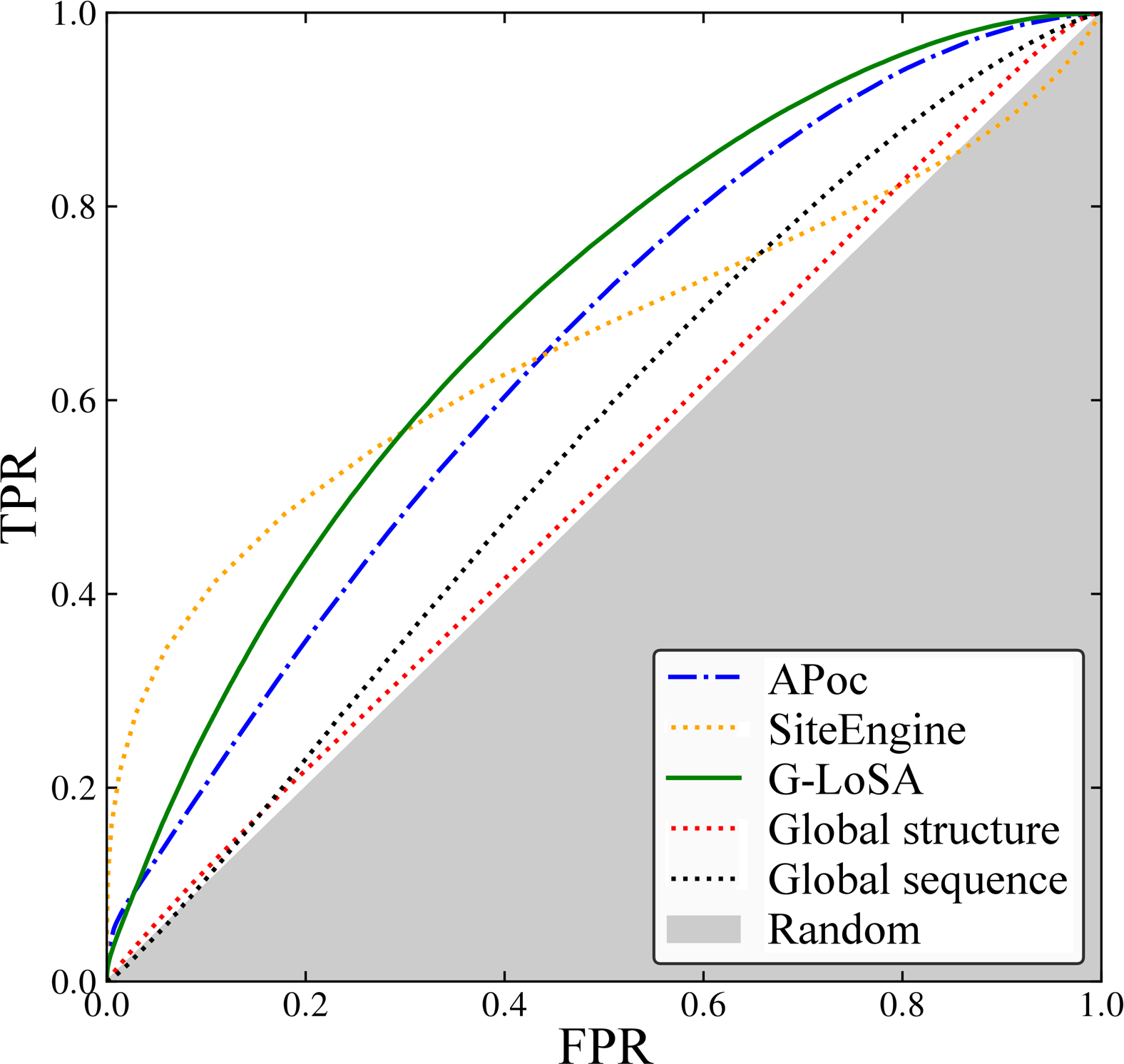
ROC plot evaluating the performance of pocket matching algorithms on the TOUGH-M1 dataset. The accuracy of APoc, SiteEngine and G-LoSA is compared to global sequence and structural alignments. The *x*-axis shows the false positive rate (FPR) and the *y*-axis shows the true positive rate (TPR). The gray area represents a random prediction.

### Characteristics of alignments constructed for the TOUGH-M1 dataset

We next investigate the correlation between the global structure similarity and the local alignment score reported by individual binding site matching algorithms for the TOUGH-M1 dataset. Figure 10 shows that local alignment scores are uncorrelated with the TM-score for APoc (Figure 10A), SiteEngine (Figure 10B), and G-LoSA (Figure 10C). These results are expected because both Positive as well as Negative pairs of proteins in TOUGH-M1 have different structures at a TM-score of <0.4. Further, binding site algorithms tested in this study have been reported to align protein binding sites in a sequence order-independent way to compute local alignment scores. On that account, similar to the APoc dataset, we check the order of alignments constructed across the TOUGH-M1 dataset with Kendall *τ*. Although alignments generated by APoc for TOUGH-M1 are notably less dependent on the sequence order compared to those constructed for the APoc dataset, the average Kendall *τ* ±standard deviation is 0.15 ±0.37 for Positive and 0.13±0.42 for Negative pairs (Figure 10D). Thus, the ordinal association of binding site residues can still be detected in APoc alignments. In contrast, the average Kendall *τ* ±standard deviation for Positive and Negative pairs is, respectively, 0.03 ±0.36 and 0.01 ±0.41 for SiteEngine (Figure 10E), and 0.06 ±0.31 and 0.07 ±0.35 for G-LoSA (Figure 10F), demonstrating that these alignments are indeed fully sequence order-independent.

**Figure 10.**
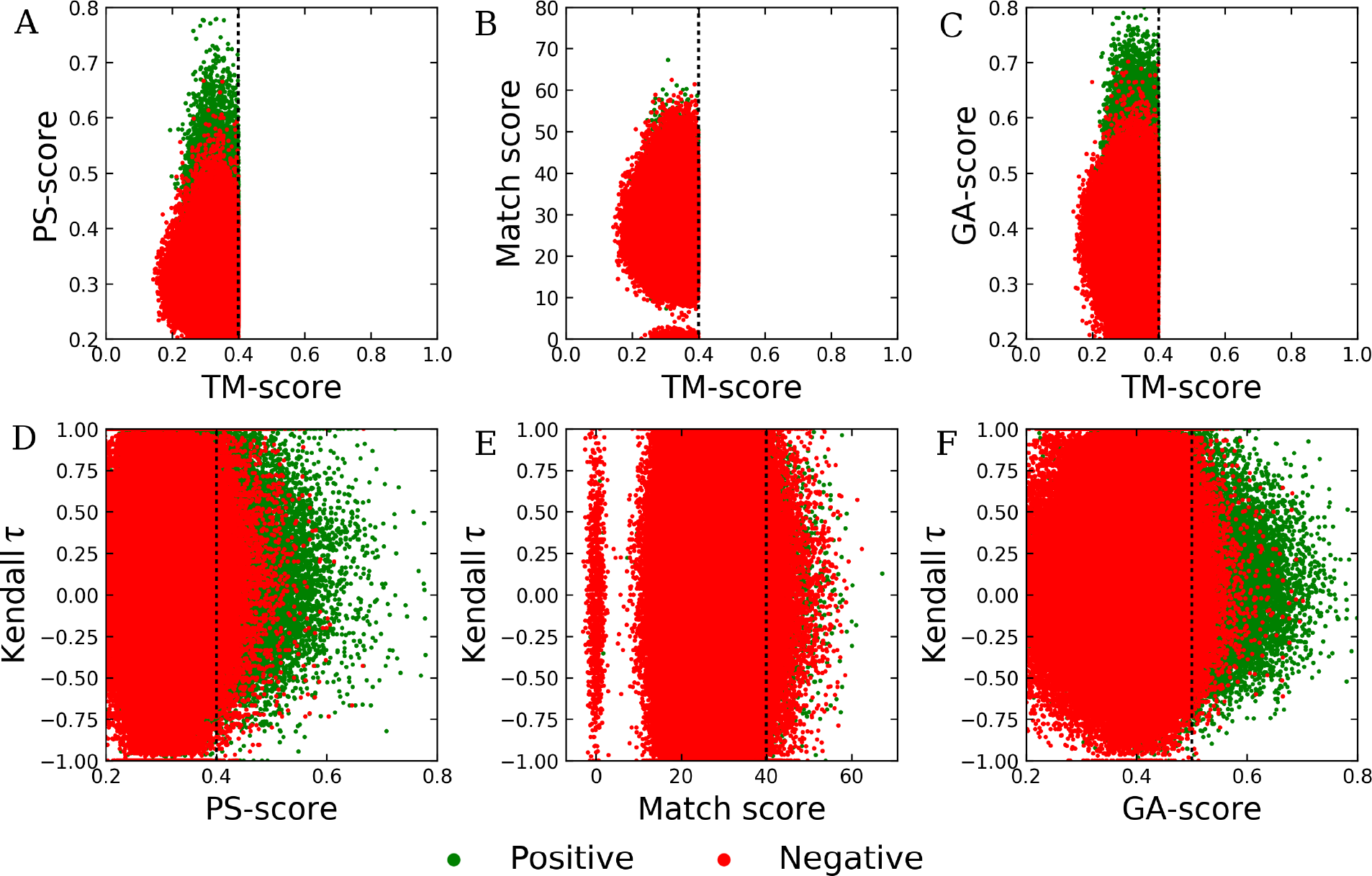
Characteristics of local alignments constructed by pocket matching algorithms for the TOUGH-M1 dataset. Top panel shows the correlation between the global structure similarity score, TM-score, and the local alignment score, (**A**) PS-score by APoc, (**B**) Match score by SiteEngine, and (**C**) GA-score by G-LoSA. Bottom panel shows the correlation between local alignment scores reported by individual pocket matching algorithms, (**D**) APoc, (**E**) SiteEngine, and (**F**) G-LoSA, and Kendall *τ* measuring the degree of the ordinal association of binding site alignments. Dotted lines mark thresholds for statistically significant alignments.

### Quantifying pocket similarity with virtual screening

Structure-based virtual screening can, in principle, be used as an indirect approach to match binding sites with the underlying assumption that similar pockets should yield similar ranking by the predicted binding affinity. To validate this approach, we first conducted a self-docking test for TOUGH-M1 targets in order to evaluate the accuracy of molecular docking with Vina and rDock. Predicted ligand-binding poses are assessed in Figure 11 by a heavy atom RMSD and the Contact Mode Score (CMS) [62] against experimental complex structures. The median RMSD for Vina is 4.07 Å, which is significantly lower than the median RMSD of 6.32 Å for rDock (Figure 11A). The recently developed CMS evaluates docked poses by calculating the overlap between intermolecular contacts in the predicted and experimental complex structures [62]. It offers certain advantages over the RMSD, for instance, the CMS is ligand size-independent providing a better evaluation metric for heterogeneous datasets, such as TOUGH-M1. Consistent with the RMSD-based assessment, Figure 11B shows that the median CMS values for Vina and rDock are 0.31 and 0.17, respectively. Our benchmarking calculations demonstrate that Vina outperforms rDock in binding pose prediction, which is in line with previous studies [41, 63].

**Figure 11.**
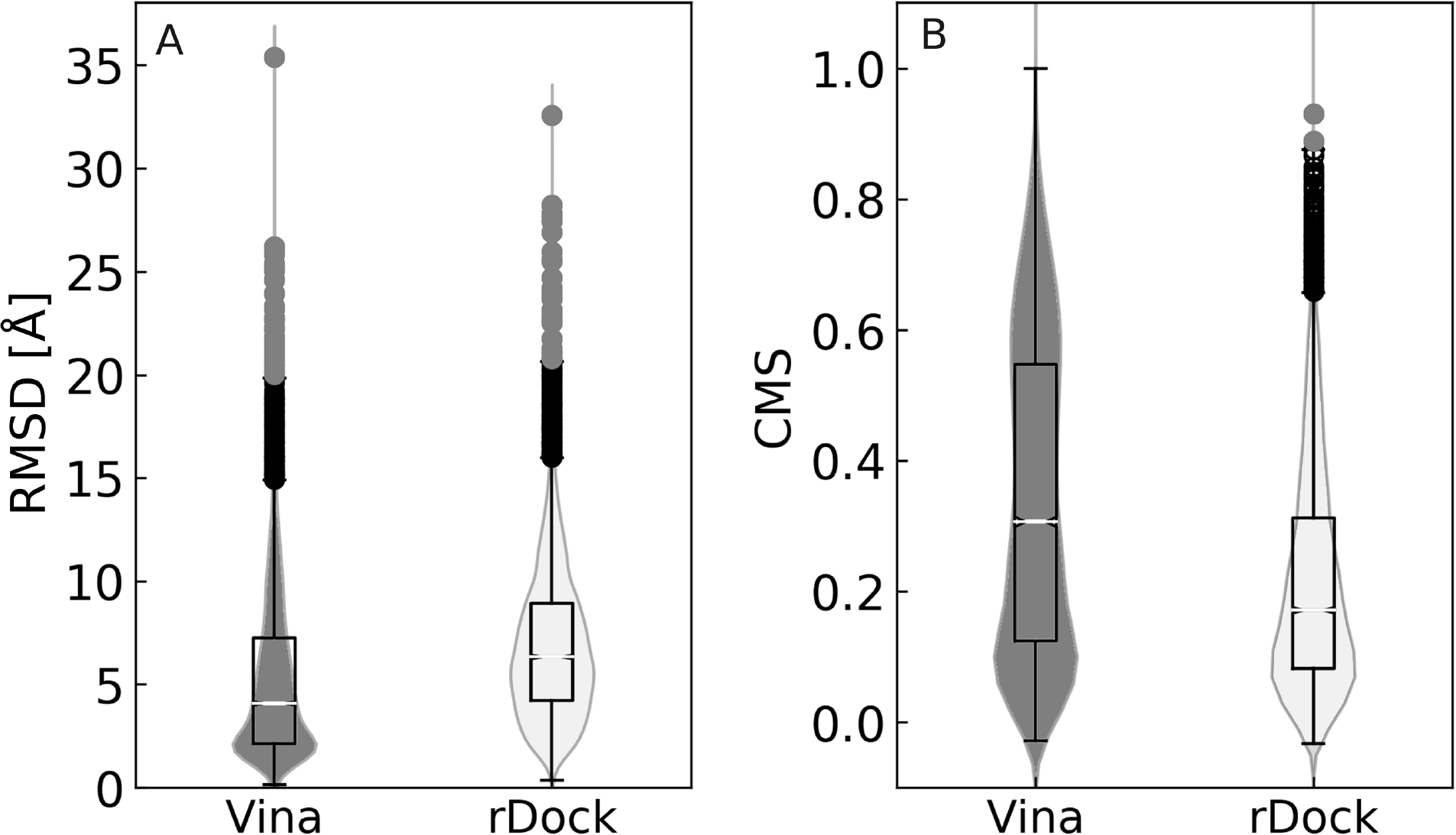
Violin plots assessing the performance of ligand docking algorithms on the TOUGH-M1 dataset. The accuracy of binding modes predicted by Vina and rDock are evaluated against experimental structures with (**A**) the root-mean-square deviation, RMSD, and (**B**) the Contact Mode Score, CMS. Boxes on the top of violins end at quartiles Q_1_ and Q_3_, and a horizontal line in each box is the median. Whiskers point at the farthest points that are within 1.5 of the interquartile range.

Subsequently, a collection of 1,515 FDA-approved drugs were docked into computationally predicted binding pockets of target proteins in the TOUGH-M1 dataset with two molecular docking algorithms, AutoDock Vina [40] and rDock [41]. Here, we test the assumption that similar pockets should produce similar ranking in structure-based virtual screening. Table 2 reports that the average Spearman’s ρ calculated for Positive pairs binding similar compounds is higher than that computed for Negative pairs binding different molecules regardless of the docking program used. Although pairs of dissimilar pockets ideally should have Spearman’s ρ around 0, these values are actually quite high, e.g. ρ is 0.67 ±0.21 for Vina. The reason for this result is that docking scores reported by many docking algorithms are highly correlated with the molecular weight of docking compounds [64-66]. On that account, we calculated Spearman’s ρ between binding affinities predicted by docking and the molecular weight of library compounds to corroborate previous studies. As expected, Table 2 shows that these values are quite high as well, e.g. ρ is 0.54 ±0.27 for Vina. For comparison, simply replacing molecular weight with random numbers yields Spearman’s ρ of around 0 corresponding to the lack of any correlation.

**Table 2.** Correlation between compound ranks from structure-based virtual screening. Vina and rDock were employed to screen a library of FDA-approved drugs against target sites in the TOUGH-M1 dataset. Average Spearman’s ρ rank correlation coefficient ±standard deviation values are reported. Positive and Negative sets correspond to pairs of TOUGH-M1 proteins binding similar and dissimilar molecules, respectively. For comparison, we include Spearman’s ρ between binding affinities calculated against each target by ligand docking algorithms, and the molecular weight and random numbers assigned to screening compounds.

| Comparison | Vina | rDock |
| --- | --- | --- |
| Positive set | 0.71 ± 0.16 | 0.56 ± 0.06 |
| Negative set | 0.67 ± 0.21 | 0.49 ± 0.12 |
| Molecular weight | 0.54 ± 0.27 | 0.28 ± 0.23 |
| Random | 0.00 ± 0.01 | 0.00 ± 0.02 |

Despite the fact that molecular docking scores are correlated with the molecular weight of screening compounds, structure-based virtual screening can be used as an indirect method to recognize those target pockets binding similar molecules. Table 1 shows that although the performance of Vina is significantly lower than all direct techniques to match binding sites and comparable to a classification based on the global sequence identity, an AUC of 0.67 for rDock is actually higher than those for APoc and SiteEngine. Since the capability of virtual screening to match ligand-binding pockets generally depends on the accuracy of a scoring function used in docking, our results are in line with previous studies reporting that rDock outperforms Vina in virtual screening [41, 67]. More importantly, an indirect comparison of pockets by means of structure-based virtual screening is methodologically orthogonal to direct techniques employing local binding site alignments, creating opportunities for novel meta-predictors, which are explored in the following section.

### Rationale for a meta-predictor

Finally, we evaluate the performance of a series of meta-predictors in distinguishing between similar and dissimilar ligand-binding pockets in the TOUGH-M1 dataset. Each meta-predictor combines a direct and an indirect method and calculates a similarity score simply by multiplying individual scores, i.e. the local alignment score reported by a binding site matching algorithm and Spearman’s ρ calculated based on the results of structure-based virtual screening. Encouragingly, Table 1 shows that combining different techniques improves the prediction accuracy over individual algorithms. For instance, integrating G-LoSA (an AUC of 0.69) with rDock (an AUC of 0. 67) into a simple meta-predictor yields the highest AUC of 0.77. Even the somewhat low performance of APoc on the TOUGH-M1 dataset (an AUC of 0.65) can be increased to 0.70 by multiplying its score by Spearman’s ρ by rDock. These results provide a solid rationale to include structure-based virtual screening conducted with an accurate molecular docking tool as part of protocols detecting similar ligand-binding sites in unrelated proteins.

### Representative examples from the TOUGH-M1 dataset

We selected two representative examples from the TOUGH-M1 dataset to discuss the results of binding site matching with APoc and SiteEngine in terms of potential improvements by including virtual screening. Proteins shown in Figure 12 are spermidine synthase (SPDS) from *Plasmodium falciparum* (PDB ID: 2pwp, chain A, 281 aa) [68], polyamine receptor SpuE from *Pseudomonas aeruginosa* (PDB ID: 3ttn, chain A, 330 aa) [69], phosphonoacetaldehyde dehydrogenase PhnY from *Sinorhizobium meliloti* (PDB ID: 4i3u, chain E, 473 aa) [70], and b-lactamase blaOXA-58 from *Acinatobacter baumanii* (PDB ID: 4y0u, chain A, 242 aa) [71]. SPDS and SpuE included in the Positive subset of TOUGH-M1 are the first example. Despite having unrelated sequences (19.5% identity) and structures (a TM-score of 0.34), these proteins bind the same ligand, spermidine. The superposition of both structures according to the local alignment constructed by APoc is presented in Figure 12A. The corresponding PS-score of 0.34 is below a threshold for the pocket similarity and the *p*-value of 0.16 is statistically insignificant, thus APoc failed to identify these proteins as binding similar ligands. The alignment constructed by APoc is partially sequential with the Kendall τ of 0.47. Moreover, the ligand RMSD for spermidine molecules upon the pocket superposition is 6.84 Å revealing inaccuracies in this local alignment.

**Figure 12.**
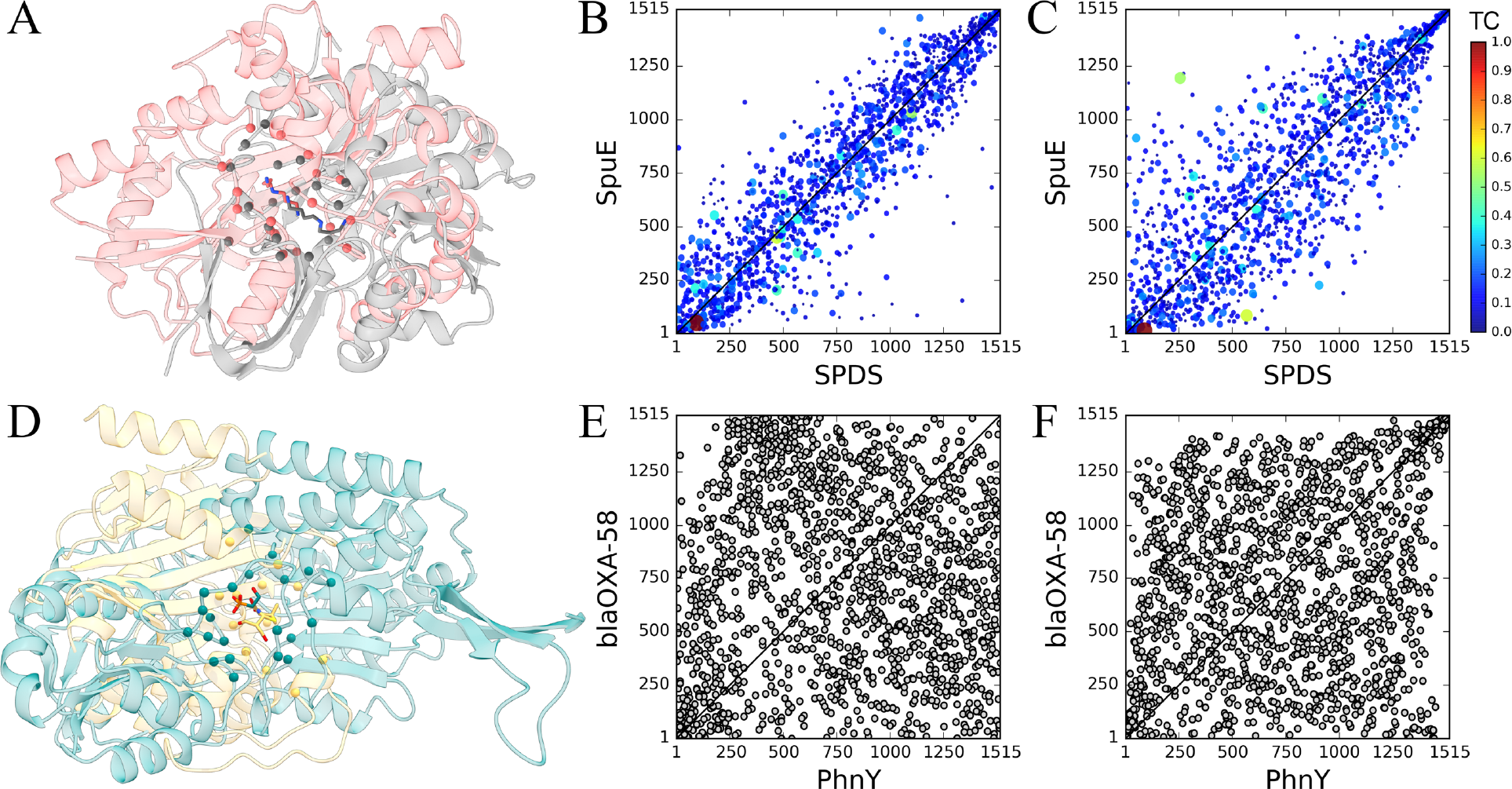
Examples of pocket alignments and the chemical correlation from structure-based virtual screening for proteins selected from the TOUGH-M1 dataset. (**A-C**) A Positive pair of spermidine synthase (SPDS) and polyamine receptor SpuE, (**D-F**) a Negative pair of phosphonoacetaldehyde dehydrogenase PhnY and b-lactamase blaOXA-58. Target structures are aligned according to local alignments reported by (**A**) APoc and (**D**) SiteEngine; SPDS, SpuE, PhnY, and blaOXA-58 are colored in salmon, gray, cyan, and yellow, respectively. Cα atoms of binding residues are represented by solid spheres, whereas binding ligands are shown as sticks. Four scatter plots show the correlation of ranks from virtual screening conducted by (**B, E**) Vina and (**C, F**) rDock. Each dot represents one library compound, whose ranks against target pockets are displayed on *x* and *y* axes. A dashed black line is the diagonal corresponding to a perfect correlation. (**B, C**) The color and size of dots depend on the Tanimoto coefficient (TC) computed against spermidine that binds to both proteins, SPDS and SpuE, in experimental complex structures; the color scale is displayed on the right.

On the other hand, results from virtual screening with Vina (Figure 12B) and rDock (Figures 12C) strongly indicate that these pockets are in fact chemically similar. Specifically, Spearman’s ρ values calculated for ranks assigned by Vina and rDock are as high as 0.91 and 0.85, respectively. This high chemical correlation is generally contingent on the accuracy of structure-based virtual screening against both target pockets, i.e. the docking program should provide reliable binding affinities to rank screening compounds. Indeed, spermidine shown as a large red dot in Figures 12B and 12C was ranked 94^th^/51^st^ against SPDS/SpuE by Vina and 89^th^/16^th^ by rDock. Further, many chemically similar compounds marked by warm colors in Figures 12B and 11C are found within the top ranks as well. Overall, despite a moderately low PS-score reported by APoc, including virtual screening certainly helps recognize SPDS and SpuE as a pair of globally unrelated proteins yet binding similar compounds.

The second example is a pair of targets, PhnY and blaOXA-58, included in the Negative subset of TOUGH-M1. These proteins have unrelated sequences (17.9% identity) and structures (a TM-score of 0.31), and bind chemically different ligands, PhnY binds phosphonoacetaldehyde and blaOXA-58 binds 6a-hydroxymethyl penicillin derivative, whose pairwise TC is only 0.16. Figure 12D shows the local alignment between PhnY and blaOXA-58 constructed by SiteEngine. This pair of targets was assigned a fairly high Match score of 45, incorrectly indicating that PhnY and blaOXA-58 bind similar compounds. Further, the pocket alignment by SiteEngine is partially sequential with the Kendall *τ* of 0.30. This obvious over-prediction by SiteEngine can be counterbalanced by virtual screening because Spearman’s ρ is as low as 0.03 for Vina (Figure 12E) and 0.27 for rDock (Figure 12F). Both docking programs produced uncorrelated ranks in virtual screening against PhnY and blaOXA-58, correctly recognizing that these targets bind chemically different molecules. Case studies presented in this section illustrate how the accuracy of direct methods to detect similar pockets in globally unrelated proteins can be enhanced by including structure-based virtual screening as an indirect component in order to reduce the number of false positives and false negatives.

## Conclusions

In this communication, we comprehensively evaluate the performance of several programs developed to identify similar binding sites in proteins. Benchmarking APoc, SiteEngine, and G-LoSA against the existing APoc dataset leads to an overestimated accuracy of these algorithms because more than one-third of similar pocket pairs come from globally similar proteins. To address this issue, we compiled TOUGH-M1, a high-quality dataset to conduct rigorous assessments of pocket matching tools. Eliminating global similarities between target proteins in the TOUGH-M1 dataset causes the performance of APoc to drop by 17% and G-LoSA by 8%, whereas the performance of SiteEngine increases by 6%. Moreover, pocket matching programs selected for this study were reported to generate sequence order-independent alignments of ligand-binding sites. Nevertheless, the analysis of the sequential ordinal association of alignments generated by these algorithms reveals that only G-LoSA produces alignments fairly independent on the sequence order. Compared to APoc and SiteEngine, G-LoSA offers a better performance detecting similar pockets across the TOUGH-M1 dataset, however, the accuracy of the constructed local alignments is somewhat unsatisfactory. The quality of pocket alignments needs to be significantly improved in order to employ ligand-binding poses predicted by G-LoSA in rational, structure-based drug repositioning.

In addition, we compare the performance of algorithms directly matching binding pockets to an indirect strategy employing structure-based virtual screening with AutoDock Vina and rDock. Although this approach, particularly with rDock as a docking program, offers a similar performance to alignment-based pocket matching techniques, combining direct and indirect methods into a meta-predictor outperforms individual algorithms. Encouragingly, the performance of pocket matching tools against the TOUGH-M1 dataset can be increased by 5% (APoc) to 10% (SiteEngine) by including virtual screening with rDock. Overall, meta-predictors employing methodologically orthogonal techniques offer a new state-of-the-art in ligand-binding site matching. This improved accuracy can beneficially be exploited in protein function inference, polypharmacology, drug repurposing, and drug toxicity prediction, accelerating the development of new biopharmaceuticals.

## Declarations

### Ethics

Not applicable.

### Consent to publish

Not applicable.

### Competing interests

The authors declare that the research was conducted in the absence of any commercial or financial relationships that could be construed as a potential conflict of interest.

### Authors’ contributions

MB designed the study, prepared datasets and developed protocols. RGG performed calculations and analyzed the data. RGG and MB wrote the manuscript. All authors read and approved the final manuscript.

### Availability of data and materials

All data reported in this paper are freely available to the community through the Open Science Framework at https://osf.io/6ngbs/.

### Funding

Research reported in this publication was supported by the National Institute of General Medical Sciences of the National Institutes of Health under Award Number R35GM119524. The content is solely the responsibility of the authors and does not necessarily represent the official views of the National Institutes of Health.

### List of abbreviations used

APoc: Alignment of Pockets
AUC: area under the curve
BLAST: Basic Local Alignment Search Tool
CMS: Contact Mode Score
DP: Dynamic Programing
FDA: U.S. Food and Drug Administration
FPR: false positive rate
G-LoSA: Graph-based Local Structure Alignment
GA-score: G-LoSA Alignment score
GSHase: glutathione synthetase
IP(3)-3KB: inositol 1,4,5-trisphosphate 3-kinase B
Ipk2: inositol polyphosphate multikinase 2
LINCL: late infantile neuronal ceroid lipofuscinosis
LPC: Ligand-Protein Contacts
MCC: Matthews correlation coefficient
PDB: Protein Data Bank
PCC: Pearson correlation coefficient
PS-score: Pocket Similarity score
RMSD: root-mean-square deviation
ROC: Receiver Operating Characteristic
SSC: Szymkiewicz-Simpson overlap coefficient
SOIPPA: Sequence Order-Independent Profile-Profile Alignment
SPDS: spermidine synthase
TC: Tanimoto coefficient
TM-score: Template Modeling score
TPP1: tripeptidyl-peptidase I
TPR: true positive rate
VS: virtual screening

